# Elevated alpha-1 antitrypsin is a major component of GlycA-associated risk for future morbidity and mortality

**DOI:** 10.1101/309138

**Authors:** Scott C. Ritchie, Johannes Kettunen, Marta Brozynska, Artika P. Nath, Aki S. Havulinna, Satu Männistö, Markus Perola, Veikko Salomaa, Mika Ala-Korpela, Gad Abraham, Peter Würtz, Michael Inouye

## Abstract

Integration of electronic health records with systems-level biomolecular data has led to the discovery that GlycA, a complex nuclear magnetic resonance (NMR) spectroscopy biomarker, predicts long-term risk of disease onset and death from myriad causes. To determine the molecular underpinnings of the disease risk of the heterogeneous GlycA signal, we used machine learning to build imputation models for GlycA’s constituent glycoproteins, then estimated glycoprotein levels in 11,861 adults across two population-based cohorts with long-term follow-up. While alpha-1-acid glycoprotein had the strongest correlation with GlycA, our analysis revealed that alpha-1 antitrypsin (AAT) was the most predictive of morbidity and mortality for the widest range of diseases, including heart failure (HR=1.60 per s.d., P=1×10^−10^), influenza and pneumonia (HR=1.37, P=6×10^−10^), and liver diseases (HR=1.81, P=1×10^−6^). Despite emerging evidence of AAT's role in suppressing inflammation, transcriptional analyses revealed elevated expression of diverse inflammatory immune pathways with elevated AAT levels, suggesting AAT is elevating to compensate for low-grade chronic inflammation. This study clarifies the molecular underpinnings of the GlycA biomarker and its associated disease risk, and indicates a previously unrecognised association between elevated AAT and severe disease onset and mortality.

## Introduction

The identification and characterisation of new predictive biomarkers for disease is fundamental to precision medicine^1,2^. Biomarkers discovered using systems-level technologies can be complex and heterogeneous, thus it can be challenging to pinpoint relevant biomolecular pathways. Therefore, knowledge of the underlying molecular basis for a biomarker is critical for identifying potential therapeutic targets and interventions.

Of recent interest is the GlycA biomarker, a serum NMR signal that has been shown to be highly predictive of morbidity and mortality from diverse diseases^3,4^, including cardiovascular diseases^5–8^, certain cancers^7–9^, type II diabetes^5,10,11^, liver diseases^5,12^, chronic inflammatory conditions^5,8^, renal failure^5^, severe infections^13^, and all-cause mortality^7,8^. Elevated GlycA levels are associated with inflammation arising from recent infection, injury, or chronic disease^14–18^, as well as low-grade chronic inflammation that may persist for up to a decade in otherwise apparently healthy adults^13^. Interestingly, the associations between elevated GlycA and disease morbidity and mortality have been largely independent of C-reactive protein (CRP)^5–13^ with suggestions that GlycA better captures systemic inflammation due to its composite nature^3,15,19^. The GlycA signal is an agglomeration of at least five circulating glycoprotein concentrations: predominantly alpha-1 antitrypsin (AAT), alpha-1-acid glycoprotein (AGP), haptoglobin (HP), transferrin (TF), and alpha-1-antichymotrypsin (AACT)^14,15^. The heterogeneous composition of GlycA represents a challenge for further research towards investigating and developing molecular intervention strategies. This is compounded by the dynamic nature of each glycoprotein, each of which responds over different times scales, directions, and magnitudes as part of the inflammatory response^13–15,20^. Thus, two individuals with the same GlycA levels may have differing concentrations of each glycoprotein contributing to the NMR spectral signal. Further, high-throughput NMR spectroscopy cannot measure the concentrations of the individual glycoproteins comprising GlycA, which require the use of specialised immunoassays. However, such immunoassays are costly and time-consuming. Here, we decompose the spectral GlycA biomarker by developing imputation models for GlycA's constituent glycoproteins, then utilise these imputed molecular phenotypes to investigate associations with disease risk. Our findings provide important insights into potential intervention strategies for GlycA-associated disease and mortality risk and may lead to better disease risk stratification.

## Results

We utilised machine learning together with matched serum NMR-metabolite measures and immunoassays for AAT, AGP, HP, and TF to develop imputation models for the concentrations of each glycoprotein. Matched NMR-metabolite measures and glycoprotein assay data were available for 626 adults from the population-based DIetary, Lifestyle, and Genetic determinants of Obesity and Metabolic syndrome 2007 study (DILGOM07)^13,21^. Lasso regression^22,23^ was used to find the optimal subset of features and corresponding weights that most accurately predicted each glycoprotein. A 10-fold cross-validation procedure was used to train each lasso regression model to reduce overfitting and estimate model accuracy (**Figure S1, Methods**). In total, 149 metabolic measurements quantified via NMR (**Table S1**) along with participant age, sex, and body mass index (BMI) were included as features to the model training procedure. The imputation models for AAT, AGP, HP, and TF explained 43%, 64%, 56% and 18% of their variation (r^2^), respectively, (**Figure 1A**) and comprised 18, 23, 27, and 9 input features, respectively (**Table S2**).

**Figure 1:**
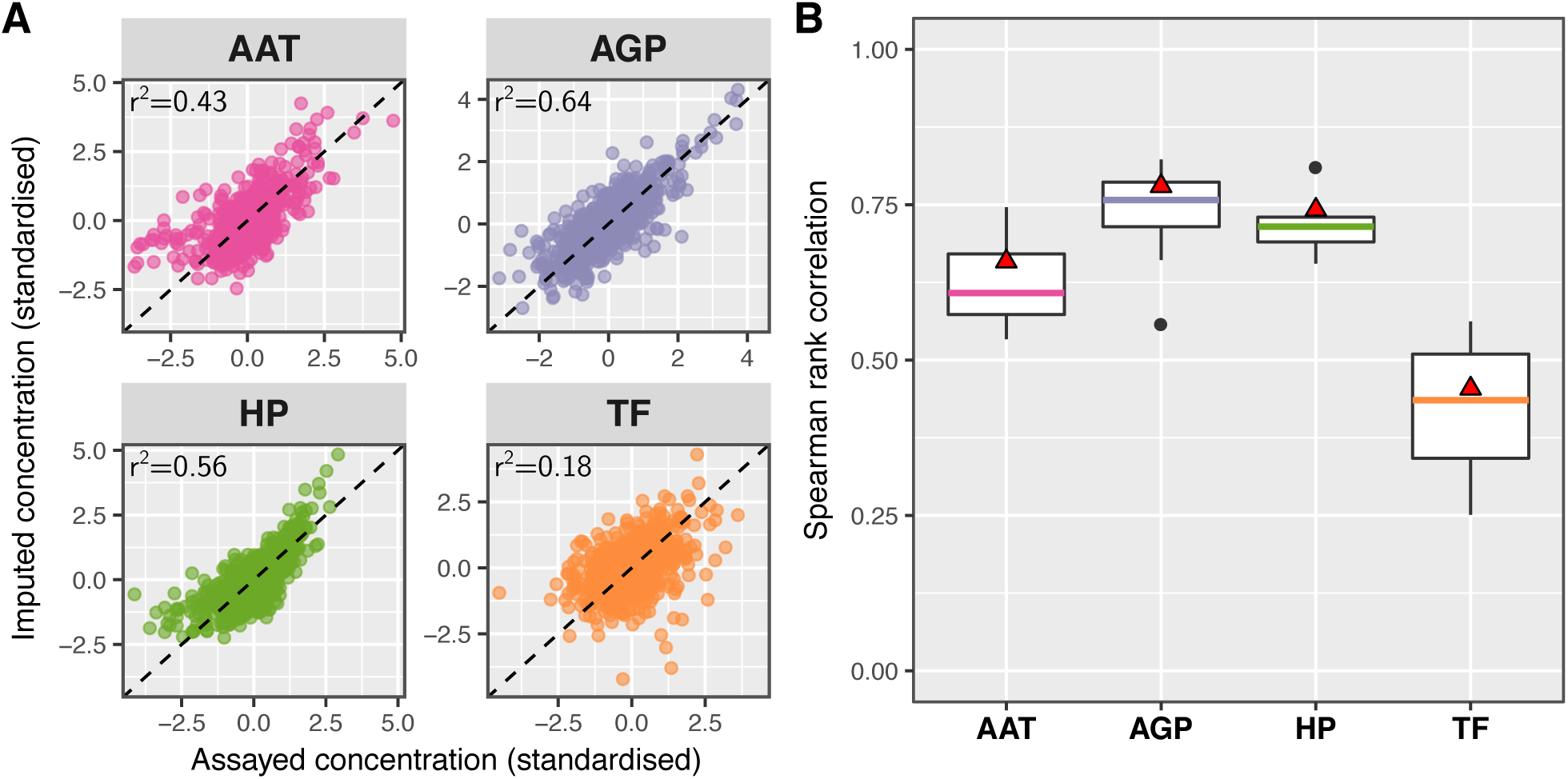
Comparison of imputation models to glycoprotein immunoassays in the 626 DILGOM07 participants with matched glycoprotein assay and metabolite quantification by NMR metabolomics. **A)** Comparison of the imputed glycoprotein levels (y-axes) to the immunoassayed glycoprotein levels (x-axes) after log transformation and standardisation. The r^2^ value indicates the proportion of variance in the assayed glycoprotein explained by the respective imputation models. **B)** Boxplots of the Spearman correlation between the imputed and observed concentrations observed in the 10-fold cross validation procedure used for model training. Red triangles show the Spearman correlation between the predicted and observed concentrations in panel **A** (detailed in **Table S3**).

Comparison of each imputation model’s predicted levels to the observed immunoassayed levels in DILGOM07 (**Figure 1A**) along with cross-validation estimates of the Spearman correlation (ρ) obtained during model training (**Figure 1B, Table S3, Methods**) indicated the imputation models for AAT (Spearman’s ρ=0.63), AGP (Spearman’s ρ=0.74), and HP (Spearman’s ρ=0.71) were sufficiently accurate for downstream analysis. In contrast, the imputation model for TF was substantially less accurate (Spearman’s ρ=0.42 and r^2^=0.18; **Figure 1, Table S3**) so was not taken forward for electronic health record association analyses.

We next imputed AAT, AGP and HP concentrations in 4,540 DILGOM07 participants and 7,321 participants from the population-based FINRISK study 1997 (FINRISK97)^24–26^, then analysed linked electronic hospital records over a matched 8-year follow-up period (**Methods**). Baseline cohort characteristics are described in **Table 1**. We observed strong, consistent, and replicable associations (FDR adjusted P<0.017, additional Bonferroni correction for the three glycoproteins) between each of AAT, AGP, and HP and increased risk of morbidity and mortality for a diverse range of disease outcomes (**Figure 2, Figure S2, Table S5**), consistent with associations seen for GlycA itself^5^. Importantly, hazard ratios calculated from the imputed measurements were consistent with those from directly assayed glycoproteins in the 630 DILGOM07 participants in which they were measured (**Figure S2**), indicating that the imputation models remained similarly accurate in the full DILGOM07 and FINRISK97 cohorts. In meta-analysis of DILGOM07 and FINRISK97, hazard ratios (HRs) were only slightly attenuated when adjusting for CRP (**Figure S3**).

**Table 1:** Cohort characteristics. Data are reported as the mean ± standard deviation (s.d.) unless otherwise indicated.

|  | **DILGOM07**<br>Model training<br>dataset | **DILGOM07**<br>Full dataset | **FINRISK97** |
| --- | --- | --- | --- |
| **Collection year** | 2007 | 2007 | 1997 |
| **Number of participants** | 626 | 4,540 | 7,321 |
| **Number (and %) of women** | 328 (53%) | 2,387 (53%) | 3,644 (50%) |
| **Mean age in years (and range)** | 53 (25–74) | 52 (25–74) | 48 (25–74) |
| **Follow-up time** | 8 years | 8 years | 8 years |
| **Body mass index (kg/m<sup>2</sup>)** | 26.80 ± 4.66 | 27.2 ± 4.8 | 26.6 ± 4.5 |
| **GlycA (mmol/L)** | 1.30 ± 0.18 | 1.30 ± 0.20 | 1.41 ± 0.25 |
| **Glycoprotein assays (# participants)** |  |  |  |
| **AAT (mg/L)** | 1.19 ± 0.20 (N=615) | 1.19 ± 0.20 (N=626) | - |
| **AGP (mg/L)** | 789 ± 203 (N=615) | 793 ± 205 (N=626) | - |
| **HP (mg/L)** | 1.09 ± 0.49 (N=614) | 1.10 ± 0.50 (N=622) | - |
| **TF (mg/L)** | 2.65 ± 0.38 (N=615) | 2.66 ± 0.38 (N=626) | - |
| **Imputed glycoproteins (# participants)** |  |  |  |
| **AAT (mg/L)** | 1.18 ± 0.11 (N=615) | 1.16 ± 0.09 (N=4,496) | 1.29 ± 0.11 (N=7,246) |
| **AGP (mg/L)** | 779 ± 145 (N=615) | 786 ± 142 (N=4,474) | 832 ± 178 (N=7,151) |
| **HP (mg/L)** | 1.04 ± 0.40 (N=614) | 1.00 ± 0.33 (N=4,491) | 1.14 ± 0.46 (N=7,194) |
| **TF (mg/L)** | 2.63 ± 0.10 (N=615) | - | - |

**Figure 2:**
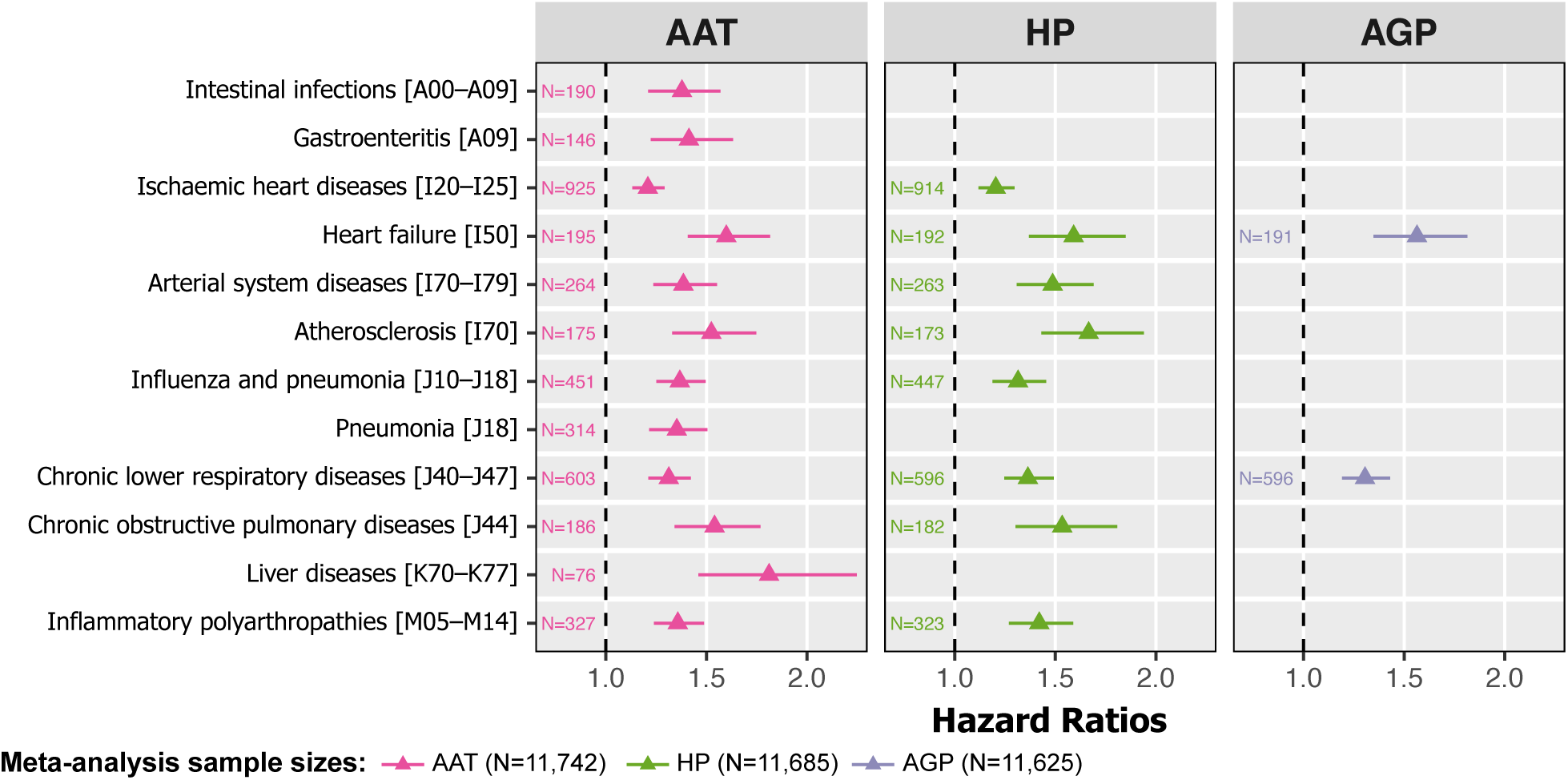
Significant and replicable glycoprotein associated risks of disease and mortality. Cox proportional hazard ratios (triangles) for the first diagnosis occurrence (hospitalisation or mortality) conferred per standard deviation increase of AAT, HP, or AGP in inverse-variance weighted fixed effects meta-analysis of DILGOM07 and FINRISK97 (**Methods**). Hazard ratios are shown only where associations were significant and replicable (Storey-Tibshirani FDR adjusted P-value < 0.05/3, adjusting for the three glycoproteins, in DILGOM07, FINRISK97, and meta-analysis). Diagnosis data were analysed for a total of 351 outcomes with >20 events in both DILGOM07 and FINRISK97 over a matched 8-year follow-up period (full listing in **Table S4**). Models were fit using age as the time scale and adjusting for sex, smoking status, BMI, blood pressure, alcohol consumption, prevalent disease prior to baseline (Methods), and previously identified biomarkers for 5-year risk of all-cause mortality (citrate, albumin, and VLDL particle size). Bars around each hazard ratio indicate the 95% confidence interval. Meta-analysis sample sizes indicate the total number of DILGOM07 and FINRISK97 participants for which the glycoprotein was successfully imputed (**Methods**). The number of events for each outcome across all participants in the meta-analysis with the corresponding imputed glycoprotein measurement are shown to the left of each hazard ratio. The alphanumeric codes in the square brackets indicate the ICD10 code or ICD10 disease group for each diagnosis. Hazard ratios fit separately in DILGOM07 and FINRISK97 along with comparison to the hazard ratios calculated from the immunoassayed AAT, HP, and AGP measurements can be found in **Figure S2** and **Table S5**. A direct comparison of meta-analysis hazard ratios across biomarkers can be found in **Figure S4** and **Table S7**.

Consistent with previous studies of GlycA^13–15^, AGP was the most strongly correlated glycoprotein with GlycA (Spearman ρ=0.65; **Table S6**). Surprisingly, despite AGP levels explaining the most variance in GlycA levels, we found that imputed AAT was significantly associated with risk of hospitalisation or death for substantially more outcomes (**Figure 2**). Elevated concentrations of imputed AAT were associated with increased 8-year risk from a wide range of disease classifications, including liver diseases (HR=1.81 per s.d. AAT, 95% CI=1.46–2.25, FDR adjusted P=1×10^−6^), heart failure (HR=1.60, 95% CI=1.41−1.82, FDR=1×10^−10^), and chronic obstructive pulmonary disease (HR=1.54, 95% CI=1.34−1.77, FDR=3×10^−8^) (full list given in **Figure 2, Table S5**). In contrast, imputed AGP was significantly associated with increased risk from only two outcomes: heart failure (HR=1.56, 95% CI=1.35−1.81, FDR=1×10^−6^) and chronic lower respiratory diseases (HR=1.31, 95% CI=1.19−1.43, FDR=2×10^−6^) (**Figure 2**).

Sensitivity analysis showed that the wide range of associations between imputed AAT and outcomes was robust to the significance threshold (**Figure 3A**). Furthermore, AAT hazard ratios tended to have the smallest standard errors across all tested outcomes (**Figure 3B**). AGP was associated with the fewest outcomes regardless of significance thresholds (**Figure 3**). Of the significant and replicable associations, HP was the strongest predictor of chronic lower respiratory diseases (HR=1.36, 95% CI=1.25−1.49, FDR=4×10^−9^), inflammatory polyarthropathies (HR=1.42, 95% CI=1.27−1.59, FDR=7×10^−8^), and atherosclerosis (HR=1.67, 95% CI=1.43−1.94, FDR=7×10^−9^) as well as the broader grouping of all arterial system diseases, while AAT was the strongest predictor for all other significant outcomes (**Figure S4, Table S7**).

**Figure 3:**
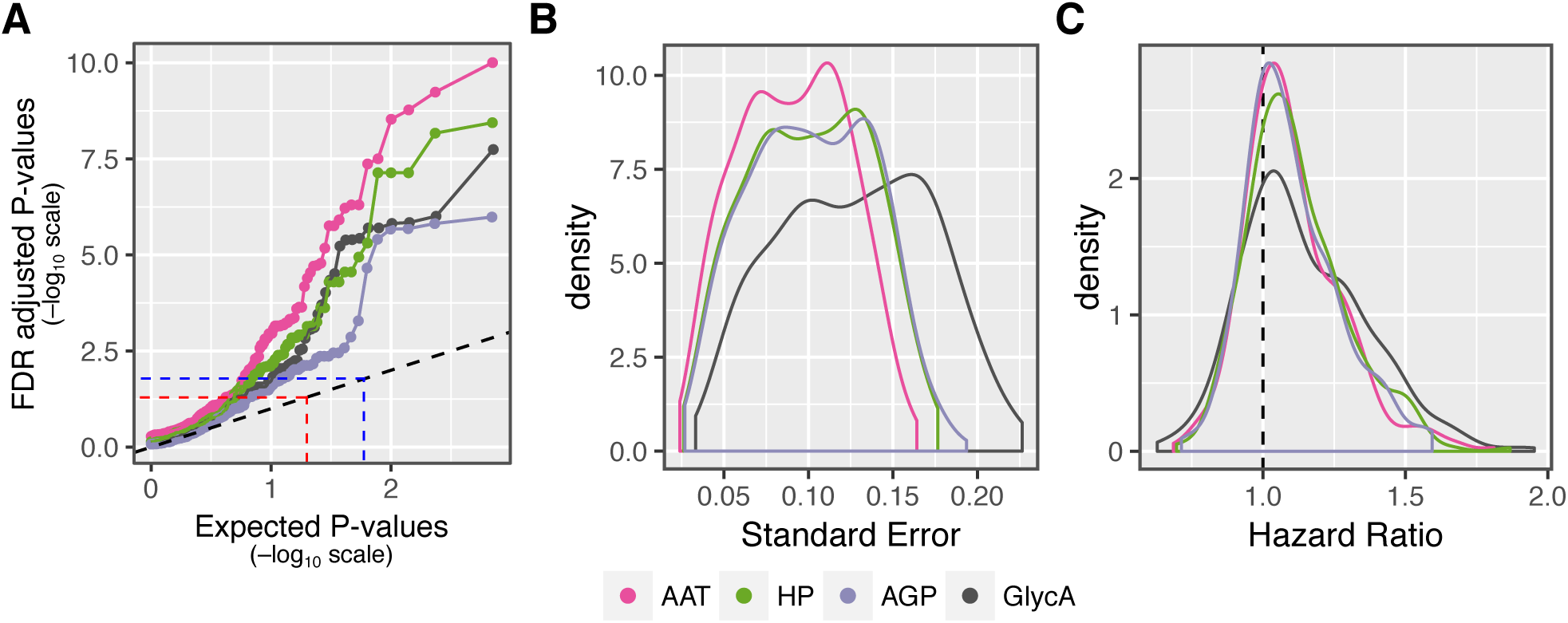
Comparison of biomarkers across all outcomes in meta-analysis of DILGOM07 and FINRISK97. **A)** Quantile-Quantile plots of distributions of hazard ratio estimate P-values (y-axis) compared to distribution of expected P-values under the null hypothesis that the corresponding biomarker is not associated with any outcome (x-axis). Hazard ratio estimate P-values are shown after adjustment for multiple testing using the Storey-Tibshirani FDR method. The dashed line indicates the location where p-values would fall if the observed distribution was identical to the null distribution. Points above the red dashed line indicated hazard ratios with FDR adjusted P < 0.05 in the meta-analysis, while points above the blue dashed line indicate hazard ratios with FDR adjusted P < 0.05/3 in the meta-analysis. Note many of the outcomes passing FDR < 0.05/3 here have FDR > 0.05/3 in either DILGOM07 or FINRISK97 thus are not included in the significant replicable outcomes shown in **Figure 2. B)** Density plots comparing each biomarker’s distribution of hazard ratio standard errors across all outcomes. **C)** Density plots comparing each biomarker’s distribution of hazard ratios across all outcomes.

While our focus here is on identifying the molecular glycoprotein associations with disease, we also performed a comparison with the GlycA NMR signal. Compared to the GlycA biomarker itself, imputed AAT was more strongly associated with a wider range of outcomes regardless of choice of significance threshold (**Figure 3A**). However, the GlycA HRs tended to be stronger than those for both AAT and HP, but with larger standard errors (**Figure 3, Figure S4**). This suggests that beyond the significant associations observed in **Figure 2**, multiple individual glycoproteins independently and weakly predict each disease, with the GlycA NMR signal capturing this risk in aggregate.

With the preponderance of AAT-associated incident disease risk and previously observed associations between GlycA and systemic inflammation^13^, we investigated whether, and to what extent, elevated AAT was associated with inflammatory processes. We used whole blood transcriptome data with matched serum AAT immunoassay data in 518 DILGOM07 participants to identify transcriptional signals in circulating immune cells associated with elevated AAT. We performed association analysis between serum AAT protein levels and 20 previously identified gene coexpression network modules^13,27–29^ to identify groups of functionally related genes associated with AAT (**Methods**) and utilised Gene Set Enrichment Analysis (GSEA)^30,31^ to identify pathways enriched for AAT-associated differential expression (**Methods**). Serum AAT protein levels were significantly associated with three gene coexpression network modules (P<0.0025; Bonferroni adjusting for the 20 tested modules) (**Table S8**). Elevated AAT corresponded to increased expression of two coexpression modules previously found to be enriched for a wide range of general immune response Gene Ontology (GO) terms^29^, and decreased expression of a module enriched for RNA processing (**Table S9**). GSEA analysis was consistent with this, finding a wide range of immune response pathways and GO terms significantly enriched for genes upregulated with elevated AAT (FDR<0.05; **Table 2, Table S10**). Elevated serum AAT protein levels were associated with increased transcription of genes involved in reactive oxygen species (FDR adjusted P=2×10^−3^), immune response initiation (*e.g*. IL6/JAK/STAT signalling, FDR adjusted P=0.02), innate immune response (*e.g*. genes localising to phagocytic vesicles, FDR adjusted P=8×10^−3^), adaptive immune response (*e.g*. Toll-like receptor signalling pathway, FDR adjusted P=0.04), and numerous cytokine regulation pathways (**Table 2, Table S10**).

**Table 2:** Highlighted gene sets significantly enriched for AAT associated genes. A selection of the gene sets that were significantly enriched for AAT-associated differential expression (**Methods**). See **Table S10** for a full listing of all 139 gene sets significantly enriched for AAT associated genes. Gene sets shown here were selected to highlight the association between elevated AAT and increase expression of diverse immune response pathways. A gene set was considered significantly enriched for AAT associated genes if its Benjamini-Hochberg FDR adjusted permutation test P-value for enrichment was < 0.05 (FDR correction performed within each gene set collection separately). The tested gene set collections included Hallmark pathways, KEGG pathways, Reactome pathways, GO biological process (GO:BP) terms, GO molecular function (GO:MF) terms, and GO cellular compartments (GO:CC). Size: number of genes on the Illumina HT-12 array annotated for the corresponding gene set. NES: enrichment score normalized by gene set size in a permutation procedure (**Methods**).

| **Collection** | **Gene set** | **Size** | **NES** | **FDR** |
| --- | --- | --- | --- | --- |
| Hallmark | Reactive oxygen species pathway | 43 | 2.21 | 0.002 |
| Hallmark | TNFα signaling via NFκB | 194 | 1.92 | 0.01 |
| Hallmark | PI3K/AKT/mTOR signaling | 101 | 1.88 | 0.02 |
| Hallmark | IL6/JAK/STAT3 signaling | 81 | 1.94 | 0.02 |
| Hallmark | Apoptosis | 154 | 1.85 | 0.02 |
| KEGG | Toll-like receptor signaling pathway | 96 | 2.05 | 0.04 |
| Reactome | Toll receptor cascades | 102 | 1.98 | 0.04 |
| GO:BP | Cytokine production involved in immune response | 17 | 2.24 | 0.006 |
| GO:BP | T cell differentiation involved in immune response | 28 | 1.96 | 0.03 |
| GO:BP | Antimicrobial humoral response | 43 | 1.97 | 0.03 |
| GO:BP | Defense response to fungus | 35 | 1.93 | 0.04 |
| GO:BP | Regulation of innate immune response | 325 | 1.86 | 0.05 |
| GO:BP | Phagocytosis engulfment | 17 | 1.87 | 0.05 |
| GO:BP | Antigen processing and presentation of peptide antigen via MHC class I | 86 | 1.86 | 0.05 |
| GO:BP | Negative regulation of viral process | 84 | 1.85 | 0.05 |
| GO:BP | Negative regulation of immune response | 113 | 1.85 | 0.05 |

## Discussion

GlycA is an NMR-based biomarker predictive of morbidity and mortality from diverse disease outcomes^3–13,19^. It is composed of the concentrations of multiple circulating glycoproteins^13–15^, each of which respond to myriad inflammatory stimuli^20^. Using circulating NMR-metabolite measures, we have developed accurate imputation models for concentrations of AAT, HP, and AGP; three of the major contributors to the GlycA signal. To investigate the molecular underpinnings of the GlycA biomarker, we imputed AAT, HP, and AGP concentrations in 11,861 generally healthy individuals from two population-based cohorts and analysed linked electronic health records over an 8-year follow-up period. Of GlycA’s constituent glycoproteins we found that alpha-1 antitrypsin, rather than alpha-1-acid glycoprotein, best explained overall future disease risk.

AAT represents a promising molecular focus for follow up studies due to its long and established history in research, well-characterised genetic variants with large effects, widely utilised diagnostic assay, and approved therapeutics. Genetic variants in AAT, such as the Z-allele, are well-known to cause AAT deficiency, which is characterised by unusually low levels of serum AAT that cause increased risk of chronic obstructive pulmonary disease/emphysema, and liver cirrhosis^32–34^. Studies have also found AAT deficiency in individuals with rheumatoid arthritis and type II diabetes^35,36^, suggesting AAT deficiency may also predispose individuals to a range of inflammation-linked disorders. Interestingly, here we found that *increased* AAT levels were predictive of morbidity and mortality for myriad common chronic diseases, suggesting that there exists a healthy window of serum AAT concentration which denotes minimal future disease risk. Robust and cost effective clinical assays measuring serum AAT concentrations are already available for diagnosis of AAT deficiency, offering a potential avenue for biomarker translation. However, further studies would be necessary to investigate AAT's role in risk prediction as well as appropriate thresholds for clinical decision making for each individual disease.

Although genetically-reduced AAT levels have been shown to be causal for disease risk, this is unlikely to be the case for elevated AAT levels. AAT is an acute-phase reactant with concentrations rising 3–4x above basal levels with inflammation due to tissue injury, infection or other exogenous insult, and may not return to normal levels for up to 6 days^20,37,38^. AAT has been found to have immunomodulating effects, and its role in regulating inflammation is being increasingly understood^39^. GlycA itself also exhibits acute-phase characteristics, and we have previously found that increased GlycA levels in population-based cohorts are associated with a low-grade inflammatory state in otherwise apparently healthy adults that likely persists for up to a decade^13^. Our transcriptional analysis showed a systemic increase in gene expression for inflammatory immune processes with elevated AAT. Since the cohort analysed here was population-based, this systemic increase in immune system activity is unlikely to reflect acute inflammation but rather is consistent with the presence of low-grade inflammation in these individuals. Taken together, increased AAT levels may reflect a response to pro-inflammatory processes.

Chronic inflammation itself contributes to the pathophysiology of common chronic diseases and development of anti-inflammatory therapies have been of interest for reducing inflammation in order to slow disease progression^40–44^. For example, recent clinical trials found an antiinflammatory Canakinumab, a monoclonal antibody for IL-1β, significantly reduced incidence of recurrent cardiovascular events as well as lung cancer in patients with previous myocardial infarction and elevated CRP^43,44^. Therapeutic administration of AAT (e.g. prolastin) is being trialled to reduce chronic inflammation for preventing the development and progression of type I diabetes, rheumatoid arthritis, and allograft rejection^45^. While we cannot make inferences about causality, our findings suggest that, if these trials are successful, an AAT therapy may have wide applicability across a range of diseases, including cardiovascular diseases. On the other hand, our results also suggest that AAT therapy may lead to increased adverse infection events as observed in the Canakinumab trial^43,44^ and dosages would need to be carefully tuned.

Our study has several limitations. Although our results suggest that alpha-1 antitrypsin is a major predictive component of the GlycA biomarker, these results are based on imputed molecular measures, and thus regression dilution (bias towards the null as measurement noise increases) may be affecting our results. However, since the imputation model for AAT had greater noise than those for AGP and HP and we observed no difference in overfitting between the three models, we do not expect that regression dilution is substantially affecting our conclusions. In addition, we cannot preclude significant associations between elevated TF or AACT and morbidity and mortality risk, for which we were unable to develop accurate imputation models. Finally, we were not able to assess causal effects of glycoproteins on incident disease.

This study demonstrates the power of machine learning for imputation of biomolecules for electronic health record-driven association analysis. Our results uncover a previously unrecognised relationship between elevated AAT, increased inflammation, and the risk of morbidity and mortality across a wide spectrum of common chronic diseases.

## Methods

### Study cohorts

In this study, we analysed data from two population-based cohorts. All cohort participants provided written informed consent. Protocols were designed and performed according to the principles of the Helsinki Declaration. Data protection, anonymity, and confidentiality have been assured. Ethics for the DILGOM07 and FINRISK97 cohort studies were approved by the Coordinating Ethical Committee of the Helsinki and Uusimaa Hospital District.

The 2007 collection of the Dietary, Lifestyle, and Genetic determinants of Obesity and Metabolic syndrome study (DILGOM07) cohort is an extension of the 2007 collection of FINRISK: a crosssectional survey of the working age population in Finland conducted every 5 years^24,25^. In DILGOM07, a detailed follow-up of 5,024 individuals was conducted to collect blood samples for omic profiling, physiological measurements, and detailed surveys of lifestyle, psycho-social, and clinical questions to study the factors leading to obesity and metabolic syndrome^21^. Serum NMR profiling was conducted for 4,816 participants; AAT, AGP, HP, and TF were measured by immunoassays for 630 participants^13^; and whole blood microarray profiling was available for 518 participants^27,28^. AACT was not available in this cohort as immunoassay measurements were performed prior to its establishment as a significant contributor to the GlycA signal by ref. 15. A total of 626 participants had matched glycoprotein assay and NMR data, and 518 participants had matched glycoprotein assay and gene expression data.

The 1997 collection of the National FINRISK study (FINRISK97) cohort contains 8,446 individuals who responded of 11,500 randomly recruited from the five major regional and metropolitan areas in Finland to monitor the health of the adult population (**aged 25–74**)^24,25^. Serum NMR profiling was conducted for 7,602 participants with adequate serum sample available^26^. Importantly, each FINRISK collection is an independent survey; the DILGOM07 and FINRISK97 cohorts are independent of one another.

### Data quantification, processing, and quality control

Venous blood samples were collected from participants in both cohorts. For DILGOM07 venous blood was drawn after an overnight fast. For FINRISK97 the median fasting time was five hours. Serum samples were subsequently aliquoted and stored at −70C.

Concentrations of circulating AAT, AGP, HP, and TF were quantified from serum samples from 630 DILGOM07 participants (626 for HP) as previously described^13^ using module analysers and Roche Tina-quant turbidimetric immunoassays. The intra-individual coefficient of variation was <3% for all four assays.

Concentrations of 228 circulating metabolites, proteins, amino acids, lipids, lipoproteins, lipoprotein subclasses and constituents, and relevant ratios were quantified by NMR metabolomics from serum samples for 4,816 DILGOM07 participants and 7,602 FINRISK97 participants. Experimental protocols including sample preparation and spectroscopy are described in ref. 46. NMR experimentation and metabolite quantification of serum samples were processed by the 2016 version of the Nightingale platform (Nightingale Health Ltd; https://nightingalehealth.com/) using a Bruker AVANCE III 500 MHz ^1^H-NMR spectrometer and proprietary biomarker quantification libraries. NMR measurements with irregular concentrations were removed and concentrations below lower detection limits set to zero by the Nightingale quality control pipeline. To facilitate log transformation, we set all zero NMR measurements to the minimum value of their respective molecular species in each cohort to approximate their lower detection limits.

Concentrations of high-sensitivity C-reactive protein (CRP) were quantified from serum samples for 4,816 DILGOM07 participants and 7,599 FINRISK97 participants using a latex turbidimetric immunoassay kit with an automated analyser.

Genome-wide gene expression profiling of whole blood for 518 DILGOM07 participants was performed as previously described^13,27,28^. Briefly, stabilised total RNA was obtained using the PAXgene Blood RNA system using the manufacturer recommended protocol. RNA integrity and quantity was evaluated for each sample using an Agilent 2100 Bioanalyser. RNA was then hybridised to Illumina HT-12 version 3 BeadChip arrays. Biotinylated cRNA preparation and BeadChip hybridisation were performed in duplicate for each sample. Microarrays were background corrected using the Illumina BeadStudio software. Probes mapping to erythrocyte globin components, non-autosomal chromosomes, or which hybridised to multiple genomic positions >10Kb apart were excluded from the analysis. Probe expression levels were obtained by taking a weighted bead-count average of their technical replicates then taking a log2 transform. Finally, expression levels for each sample were quantile normalised.

### Imputation model training

Lasso regression models were fit in the DILGOM07 participants to determine the contributions of the NMR-based biomarkers, participant age, sex, and BMI that best predicted the concentrations of each glycoprotein. Samples with any missing NMR data (N=11, 1.8%) were excluded. Consequently, all derived ratios in the NMR data were excluded from the analysis due to increased missingness arising from low concentration measurements in their numerator or denominator. In total, 149 NMR measurements (**Table S1**) were included in each lasso regression. In total, 615 individuals had matched glycoprotein and completed NMR metabolite data (N=611 for HP). Age was standardised, and each glycoprotein, NMR-metabolite measure, and BMI were log transformed and standardised when fitting the lasso regression models. The models were fit using *glmnet*^47^ version 2.0-2 in *R* version 3.1.3.

To reduce overfitting of the models to the 615 DILGOM07 participants (hereby “training cohort”), a 10-fold cross-validation procedure was used to tune the lasso regression λ penalty, which determined how many variables were included in the final imputation models for each glycoprotein (**Figure S1**). In this procedure, the training cohort was randomly split into 10 groups, and a sequence of 100 λ values was generated by *glmnet*. For each of these 100 λs a lasso regression was fit to each possible 9/10^ths^ of the data and the resulting model used to predict the glycoprotein concentration in the remaining 1/10^th^ of the data. To compare the accuracy of the model fit by each λ, the mean-square error (MSE) was calculated as the mean of squared difference between the predicted and observed glycoprotein in each test-fold (**Figure S1**). To obtain the final imputation models (**Table S2**) a lasso regression model was fit to the NMR-metabolite measures, age, sex, and BMI, for the full training cohort using the largest λ penalty with an average MSE within one standard error of the smallest average MSE in the crossvalidation procedure. This λ was selected as it produced the simplest possible model for each glycoprotein with a comparable average MSE to the smallest average MSE given the uncertainty in the average MSE estimate, thus further reducing model overfitting^47^.

To evaluate imputation model accuracy, the Spearman’s rank correlation coefficient (hereby Spearman correlation) was used to quantify the similarity of the imputed and immunoassayed levels of each glycoprotein (**Figure 1B, Table S3**). The Spearman correlation provides an indicator of how well the imputation models are likely to distinguish between many individuals with different glycoprotein concentrations after the standard statistical treatment of normalisation and standardisation when imputing each glycoprotein in another dataset. Estimates of the Spearman correlation given in the text were obtained by taking their averages across the 10-fold cross-validation procedure in which the Spearman correlation were calculated by comparing the imputed and immunoassayed glycoprotein levels in each 1/10^th^ of the data (shown by the boxplots in **Figure 1B**). The Spearman correlation was also calculated between the imputed and immunoassayed glycoprotein levels shown in **Figure 1A** after using the final imputation models to predict the concentration of each glycoprotein in all 626 DILGOM07 participants with serum NMR and matched glycoprotein assay data (point estimates shown in **Figure 1B**). The difference between this point estimate and the average Spearman correlation in model training (**Figure 1B, Table S3**) indicates the amount of overfitting of each model to the training cohort.

The strong correlation structure in the NMR-metabolite measurements meant that the imputation models in **Table S2** were not necessarily unique. Re-running the model training procedure lead to imputation models comprising different features but with similar accuracy to that shown in **Figure 1** and similar hazard ratio estimates as shown in **Figure S2**.

### Electronic health record analysis

Electronic health records were obtained and collated for individuals participating in the DILGOM07 and FINRISK97 studies as described in ref. 5. Briefly, electronic health records were obtained from the Finnish National Hospital Discharge Register and the Finnish National Causes-of-Death Register for individuals in DILGOM07 and FINRISK97 from 1987–2015. Electronic health records from 1987–1995 were encoded according to the International Classification of Diseases (ICD) 9^th^ revision (ICD-9) format, and converted to the 10^th^ revision format (ICD10) to match the encoding of records from 1996–2015 using the scheme provided by the Diagnosis Code Set General Equivalence Mappings from the Center for Disease Control in the United States of America (ftp://ftp.cdc.gov/pub/Health_Statistics/NCHS/Publications/ICD10CM/2011/), and were verified using the National Data Policy Group mapping scheme from the New Zealand Ministry of Health (http://www.health.govt.nz/system/files/documents/pages/masterf4.xls). Diagnoses with a mismatch of the first 3 digits in the ICD10 code between the two conversion protocols were verified manually.

Electronic health records were aggregated into distinct disease outcomes for each individual, each comprising an ICD10 disease grouping or ICD10 code at three-digit accuracy. Records were aggregated into incident and prevalent cases for each outcome for each individual. Incident cases comprised the first event (either hospital discharge diagnosis or mortality) in an 8-year follow-up from cohort baseline, chosen to match the maximum follow-up time for DILGOM07. Prevalent cases indicated whether an individual had any event for that outcome from 1987 to baseline (20 years for DILGOM07 and 10 years for FINRISK97), the maximum retrospective period available for the analysis. Main and side diagnoses were treated equally when aggregating electronic health records into incident and prevalent cases of each outcome.

Imputed AAT, imputed AGP, imputed HP, and GlycA were separately tested as biomarkers for incidence of each outcome in 4,540 DILGOM07 participants and 7,321 FINRISK97 participants with all model covariates and excluding pregnant women over the 8-year follow-up, and meta-analysed with an inverse-variance weighted fixed-effects model using the *metafor* R package^48^ version 2.0.0 (**Figure 2, Figure 3, Figures S2–S4, Table S5,S7**). Imputation of AAT was successful for 4,496 DILGOM07 participants and 7,246 FINRISK97 participants. Imputation of AGP was successful for 4,474 DILGOM07 participants and 7,151 FINRISK97 participants. Imputation of HP was successful for 4,491 DILGOM07 participants and 7,194 FINRISK97 participants. Any imputed glycoprotein measurements that were outside the range of measurements observed in the glycoprotein assays were excluded (0.64–2.58 mg/L for AAT, 362–1,880 mg/L for AGP, and 0.14–3.95 mg/L for HP), and were not imputed for participants where any of the imputation model inputs were missing. Cox proportional hazards models were fit using age as the time scale and adjusting for sex, smoking status, BMI, systolic blood pressure, alcohol consumption, and prevalent disease, as well as citrate, albumin, and VLDL particle size, which were previously identified as biomarkers for 5-year risk of all-cause mortality alongside GlycA levels in FINRISK97^7^. Each imputed glycoprotein, GlycA, albumin, citrate, BMI, systolic blood pressure, alcohol consumption and the diameter of VLDL particles were log transformed, and standardised (s.d.=1) in the statistical analyses while current smoking and sex were coded as categorical covariates. Association analyses were performed for all outcomes with ≥ 20 incident cases in both DILGOM07 and FINRISK97 in the subsets of individuals with successfully imputed concentrations of each glycoprotein. Adjustment for prevalent cases was performed where there were ≥ 10 prevalent cases in the respective subsets of individuals prior to baseline. Hazard Ratios were similar when excluding prevalent cases (**Figure S5**). In total, AAT, AGP, HP, and GlycA were tested as biomarkers for 351, 347, 350, and 356 outcomes, respectively (**Table S4**).

To control for the many related and unrelated hypothesis tests, P-values were adjusted across all outcomes for each biomarker and cohort separately using the Storey-Tibshirani positive False Discovery Rate method^49^ using the *qvalue* package version 2.4.2 in *R* version 3.2.3. This method is designed to control for multiple correlated tests such as the nested diagnoses and diagnosis categories tested in this study. We considered any glycoprotein–outcome association to be significant and replicable where its FDR adjusted P-value was < 0.05/3 (Bonferroni correcting the significance threshold of 0.05 for the three glycoproteins) in DILGOM07, FINRISK97, and in the meta-analysis (**Figure 2, Figure S2, Table S5**).

Sensitivity analysis to CRP was performed by fitting Cox proportional hazard models with CRP as an additional covariate (**Figure S3**). Hazard ratios were combined in inverse-variance weighted meta-analysis. Sensitivity analysis to prevalent disease adjustment was performed by fitting Cox proportional hazards models in the subset of individuals without any prevalent cases of each outcome using the same model parameters and covariates as described above (**Figure S5**).

To assess consistency of hazard ratios calculated from the imputed glycoproteins with those from the immunoassayed glycoproteins (**Figure S2**), Cox proportional models were fit for all DILGOM07 participants with immunoassayed glycoproteins (N=630 for AAT and AGP, N=626 for HP), and also for the predicted glycoprotein concentrations in the 615 DILGOM07 participants used to train the imputation models. In each case, analyses were restricted to the 46 outcomes with 20 or more events in the respective subsets of DILGOM07.

### Gene expression analysis

To identify functionally related gene sets in whole blood associated with AAT, linear regression models were fit between immunoassayed AAT levels and summary expression profiles for 20 replicable gene coexpression network modules that we previously identified in DILGOM07^13,27–29^ (**Table S8**). In DILGOM07, 518 participants had matched AAT immunoassay data and transcriptome-wide gene expression profiling. Regression models were adjusted for age and sex. An association between a module and AAT was considered significant where P<0.0025 (Bonferroni adjusted significance threshold for the 20 tested coexpression modules).

Identification and characterisation of gene coexpression network modules in DILGOM07 is described in ref. 29. Briefly, weighted gene coexpression network analysis (WGCNA)^50^ was used to identify clusters of genes whose expression levels were tightly correlated. Transcriptome-wide expression levels were adjusted for age and sex, then the Spearman correlation between expression levels was calculated for each pair of probes. The adjacency matrix encoding the network edge weight between each pair of probes was obtained by taking the absolute value of the Spearman correlation and exponentiating to the 5^th^ power, which was chosen via the scale-free topology criterion. Probes were subsequently hierarchically clustered based on the dis-similarity of their topological overlap, a metric which quantifies the similarity of two probes based on the strength of their shared network edge weight along with the similarity of their network edge weights to all other probes in the network. A total of 40 modules were identified using a dynamic tree cut algorithm with a minimum cluster size of 10. Summary expression profiles for each module were calculated as the eigenvector of the first principal component of each module’s expression matrix. Modules were assessed for topological replication (i.e. preservation of gene–gene connection patterns and coexpression density) in an independent population cohort of 1,650 individuals using the R package NetRep^51^, which performs permutation tests on seven module preservation statistics; 20,000 permutations were performed. Of the 40 coexpression network modules, 20 were found to be replicable (all seven module preservation statistics had a permutation test P<1×10^−3^).

Characterisation of the biological function of the RNA processing module, an AAT-associated module not yet characterised elsewhere, was performed as previously described^29^. First, a core set of genes for the module was defined through a permutation test of module membership. Each probe’s correlation with the module’s summary expression profile was compared to a null distribution of membership scores obtained by calculating the correlation between the module’s summary expression profile and the 33,691 probes in the transcriptome that did not cluster into the RNA processing module. The membership permutation test p-values were Benjamini-Hochberg FDR adjusted across 1,734 probes in the RNA processing module, and probes with FDR adjusted P<0.05 were considered core module probes robust to the WGCNA clustering parameters. Overrepresentation analysis of Gene Ontology (GO) biological process terms^52,53^ in the RNA processing module gene set was performed using GOrilla^54^ (**Table S9A**). The module was considered to be significantly enriched for any GO term where its hypergeometric test Benjamini-Hochberg FDR adjusted P<0.05. REVIGO^55^ was used to measure the semantic similarity of significantly enriched GO terms to rank terms by semantic uniqueness and dispensability (redundancy) (**Table S9B**).

To identify pathways associated with AAT levels we used GSEA^30,31^ (Java application version 2.2.4) to identify pathways enriched for genes differentially expressed with respect to AAT levels in DILGOM07. We tested enrichment for AAT-associated differential expression in collections of curated gene sets available from the Molecular Signatures Database (MSigDB) (http://software.broadinstitute.org/gsea/msigdb/collections.jsp, accessed May 25^th^ 2017). Specifically, we tested enrichment in the MSigDB Hallmark gene sets^56^; GO biological process, molecular function, and cellular compartment ontologies^52,53^; Kyoto Encyclopedia of Genes and Genomes (KEGG) pathways^57^; and Reactome pathways^58^. Gene sets were tested for enrichment in each collection separately. The Pearson correlation metric was used within GSEA to rank genes by their association with AAT. Age and sex adjusted probe expression levels and age- and sex-adjusted log-transformed AAT levels were provided as input since GSEA does not allow for adjustment of covariates. Expression levels for genes with multiple probes were obtained by taking the highest probe expression in each sample (performed by the GSEA software). After collapsing multiple probes, there were 30,281 genes in total. GSEA calculated an enrichment score for each gene set by taking the maximum of a running-sum statistic^30,31^. This running-sum statistic was calculated by iterating through all genes in descending order by AAT correlation, incrementing the running-sum statistic by a gene’s correlation with AAT if it appears in the gene set of interest, and decrementing the running-sum statistic by 1/30,281 otherwise. Normalised enrichment scores and enrichment score p-values were obtained through a permutation test procedure^30,31^ in which samples were shuffled 1,000 times. Normalised enrichment scores were calculated as the enrichment score divided by the average enrichment score for the corresponding gene set across the 1,000 permutations. Permutation test P-values were Benjamini-Hochberg FDR adjusted for multiple testing in each gene set collection separately. We considered any gene set to be significantly enriched for genes either up- or down-regulated with respect to increasing AAT levels where the enrichment FDR adjusted P<0.05 (**Table S10**).

### Data availability

The imputation models for AAT, AGP, HP are available in an R package, *imputegp* https://github.com/InouyeLab/imputegp.

## Acknowledgements

This study was supported by funding from National Health and Medical Research Council (NHMRC) grant APP1062227 and by the Victorian Government’s Operational Infrastructure Support (OIS) program. M.I. was supported by an NHMRC and Australian Heart Foundation Career Development Fellowship (no. 1061435). S.R. was supported by an Australian Postgraduate Award. G.A. was supported by an NHMRC Early Career Fellowship (no. 1090462). JK and PW were funded by Academy of Finland (grant numbers 297338 and 307247, 312476, and 312477) and Novo Nordisk Foundation (NNF17OC0026062 and 15998). V.S. was supported by the Finnish Foundation for Cardiovascular Research. M.A.K was supported by the Sigrid Juselius Foundation, Finland. MAK works in a Unit that is supported by the University of Bristol and UK Medical Research Council (MC_UU_12013/1). Thanks to Dr Jimmy Peters for his helpful feedback on the manuscript.

